# Engineered ACE2 counteracts vaccine-evading SARS-CoV-2 Omicron variant

**DOI:** 10.1101/2021.12.22.473804

**Authors:** Nariko Ikemura, Shunta Taminishi, Tohru Inaba, Takao Arimori, Daisuke Motooka, Kazutaka Katoh, Yuhei Kirita, Yusuke Higuchi, Songling Li, Tatsuya Suzuki, Yumi Itoh, Yuki Ozaki, Shota Nakamura, Satoaki Matoba, Daron M Standley, Toru Okamoto, Junichi Takagi, Atsushi Hoshino

**Author notes:** These authors contributed equally to this work (Nariko Ikemura, Shunta Taminishi, Tohru Inaba, Takao Arimori, Daisuke Motooka, Kazutaka Katoh). Corresponding author. (A.H.); (J.T.); (T.O.); (D.M.S.).

## Abstract

The novel SARS-CoV-2 variant, Omicron (B.1.1.529) contains an unusually high number of mutations (>30) in the spike protein, raising concerns of escape from vaccines, convalescent sera and therapeutic drugs. Here we analyze the alteration of neutralizing titer with Omicron pseudovirus. Sera obtained 3 months after double BNT162b2 vaccination exhibit approximately 18-fold lower neutralization titers against Omicron than parental virus. Convalescent sera from Alpha and Delta patients allow similar levels of breakthrough by Omicron. Domain-wise analysis using chimeric spike revealed that this efficient evasion was primarily achieved by mutations clustered in the receptor-binding domain, but that multiple mutations in the N-terminal domain contributed as well. Omicron escapes a therapeutic cocktail of imdevimab and casirivimab, whereas sotrovimab, which targets a conserved region to avoid viral mutation, remains effective. The ACE2 decoy is another virus-neutralizing drug modality that is free, at least in theory, from complete escape. Deep mutational analysis demonstrated that, indeed, engineered ACE2 prevented escape for each single-residue mutation in the receptor-binding domain, similar to immunized sera. Engineered ACE2 neutralized Omicron comparable to Wuhan and also showed a therapeutic effect against Omicron infection in hamsters and human ACE2 transgenic mice. Like previous SARS-CoV-2 variants, some sarbecoviruses showed high sensitivity against engineered ACE2, confirming the therapeutic value against diverse variants, including those that are yet to emerge.

**One Sentence Summary:** Omicron, carrying ∼30 mutations in the spike, exhibits effective immune evasion but remains highly susceptible to blockade by engineered ACE2.

## INTRODUCTION

The novel variant B.1.1.529 was detected in Botswana on November 11^th^, 2021 and spread rapidly and globally. On November 26^th^, the World Health Organization (WHO) classified B.1.1.529 as the Omicron variant of concern (VOC). Omicron possesses 26 to 32 mutations, 3 deletions and one insertion in the spike protein. Among these, 15 mutations are located in the receptor binding domain (RBD). Spike mutations have the potential to enhance transmissibility, enable immune evasion, or both(*1*). Compared to previous variants, Omicron contains far more mutations in the spike, and such mutations are expected to dramatically alter the characteristics of SARS-CoV-2. According to routine surveillance data from South Africa, Omicron has higher transmission and risk of reinfection due to immune evasion. In addition, multiple mutations in the RBD have been reported to impact escape from therapeutic monoclonal antibodies, even in cocktail regimen (*2-4*).

We previously developed an engineered ACE2 containing mutations to enhance affinity toward SARS-CoV-2 spike that showed virus-neutralizing capacity comparable to therapeutic monoclonal antibodies (*5*). The advantage of the ACE2-based decoy is its resistance to virus escape mutations. Mutant spikes escaping from ACE2 decoy may appear, but they would have limited binding affinity toward the native ACE2 receptors on host cells, making such resultant viruses unfit to propagate due to greatly reduced or even absent infectivity. In fact, engineered ACE2 successfully neutralized previous variant viruses as well as SARS-CoV-1 and showed no signs of vulnerability to escape mutants when added at suboptimal concentration during long-term culture (*5*).

In this report, we examined the antigenic alteration of Omicron and demonstrate that Omicron does indeed evade neutralization by vaccinated and convalescent sera. The ability of Omicron to escape was primarily due to mutations in the RBD, although those in the N-terminal domain (NTD) also contributed to some extent. A broad range of antibodies failed to neutralize Omicron, including a cocktail of imdevimab and casirivimab. However, the ACE2 decoy remained effective to Omicron as well as to some other sarbecoviruses.

## RESULTS

### Genetic and epidemic characteristics of Omicron

More than 30 amino acids were identified and compared to other variants of concern (VOC) reported in the GISAID database. We noticed that most mutations found in the Omicron strain were present in previous VOCs as a minor population (Fig. 1A). About 70% of mutations are located in the S1 subunit harboring the NTD and RBD. The NTD contains 4 missense mutations, 3 deletions, and 1 insertion, and most of these changes reside in the flexible loop region that is known to be the target of NTD-directed neutralizing antibodies (*6*). In the RBD, 10 of 15 missense mutations cluster in the receptor binding motif (RBM) (Fig. 1B). These characteristics suggest that mutations in Omicron are likely to influence the binding affinity of neutralizing antibodies, host receptors such ACE2, or both. The time course of dissemination of Omicron in the world has shown that the Delta strain was rapidly exchanged with the Omicron strain in South Africa (Fig. 1C). These data support the notion that mutations in the Omicron spike protein enhance the infection rates.

**Fig. 1.**
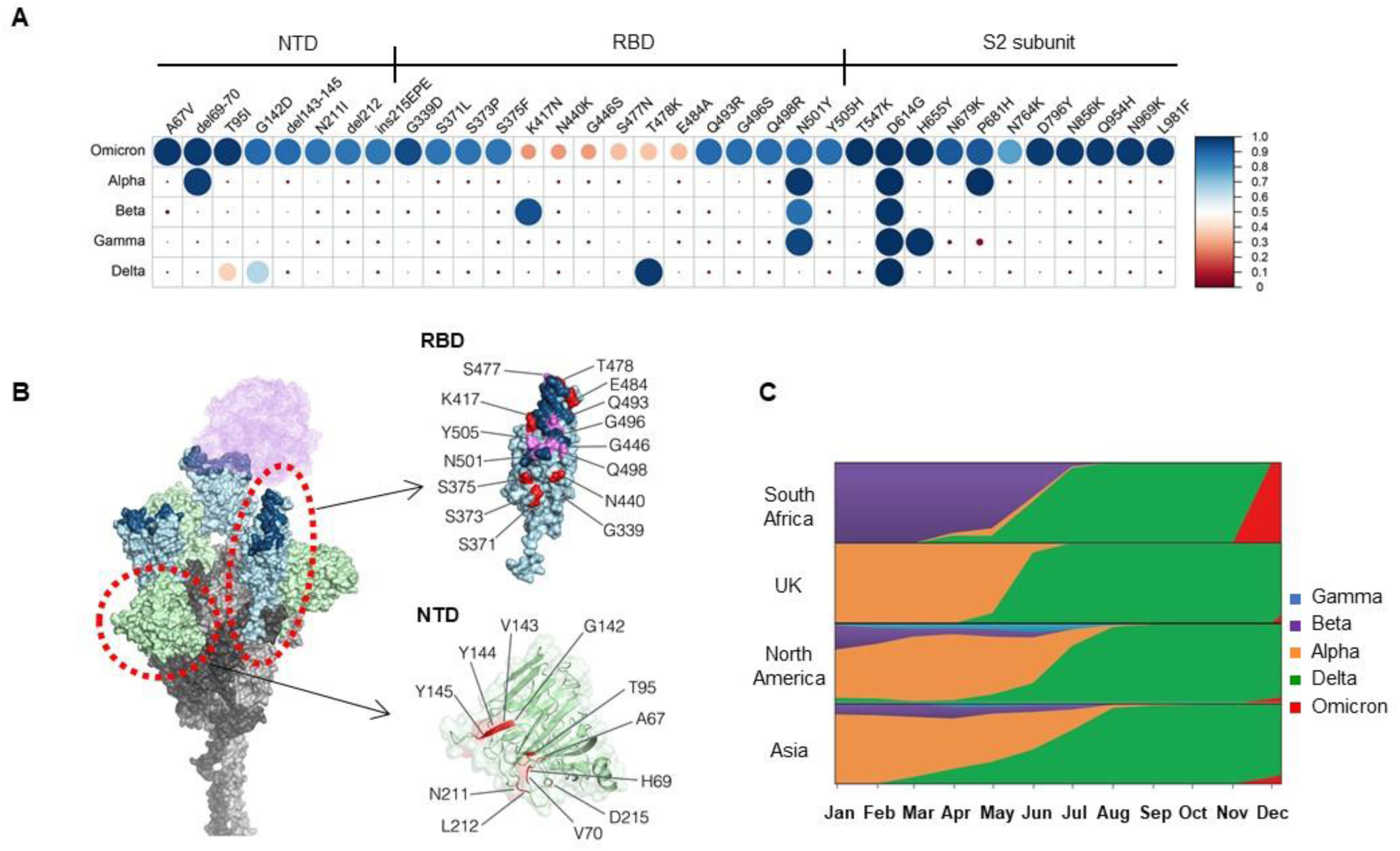
Emergence of Omicron (B.1.1.529) (**A**) Amino Acid mutations of S protein between SARS-CoV-2 strains. Mutation frequency of each amino acid substitution in spike protein which was observed in Omicron were calculated using SARS-CoV-2 sequences reported in GISAID as 17^th^ Dec 2021 (Alpha = 1,138,704, Beta = 40,135, Gamma = 117,200, Delta = 3,441,137, and Omicron = 5,469 sequences). (**B**) A structural model of SARS-CoV-2 S trimer with all three RBDs in the open state based on PDB: 7A89. In the RBD, RBM is highlighted in dark blue and Omicron mutations are highlighted in purple or red in or outside the RBM. (**C**) Relative proportion of variant cases during 2021 in South Africa, United Kingdom, North America, and Asia.

### Omicron evades vaccine and convalescent sera through mutations in the RBD and NTD

To evaluate the infectivity of Omicron, we generated a pseudotyped virus harboring the spike of Omicron and assessed neutralizing activity of BNT162b2 vaccinated and convalescent sera against it. Virus neutralization assays with vaccinated sera obtained from 12 persons 3 months after vaccination with double Pfizer-BioNTech vaccine BNT162b2 showed 17.7-fold lower neutralization titers against Omicron than the D614G mutation of the parental virus (Fig. 2A and Fig. S1). We also collected convalescent sera before or after the pandemic of the Delta variant. Convalescent sera before the Delta pandemic (December 2020 through January 2021) showed 19.3 and 17.8-fold reduction compared with parental virus or Alpha. On the other hand, sera collected during/after the Delta pandemic (August 2021 through October 2021) exhibited 9.5 and 15.4-fold reduction compared with the parental virus or Delta, respectively. (Fig. 2B, C and Fig. S1).

**Fig. 2.**
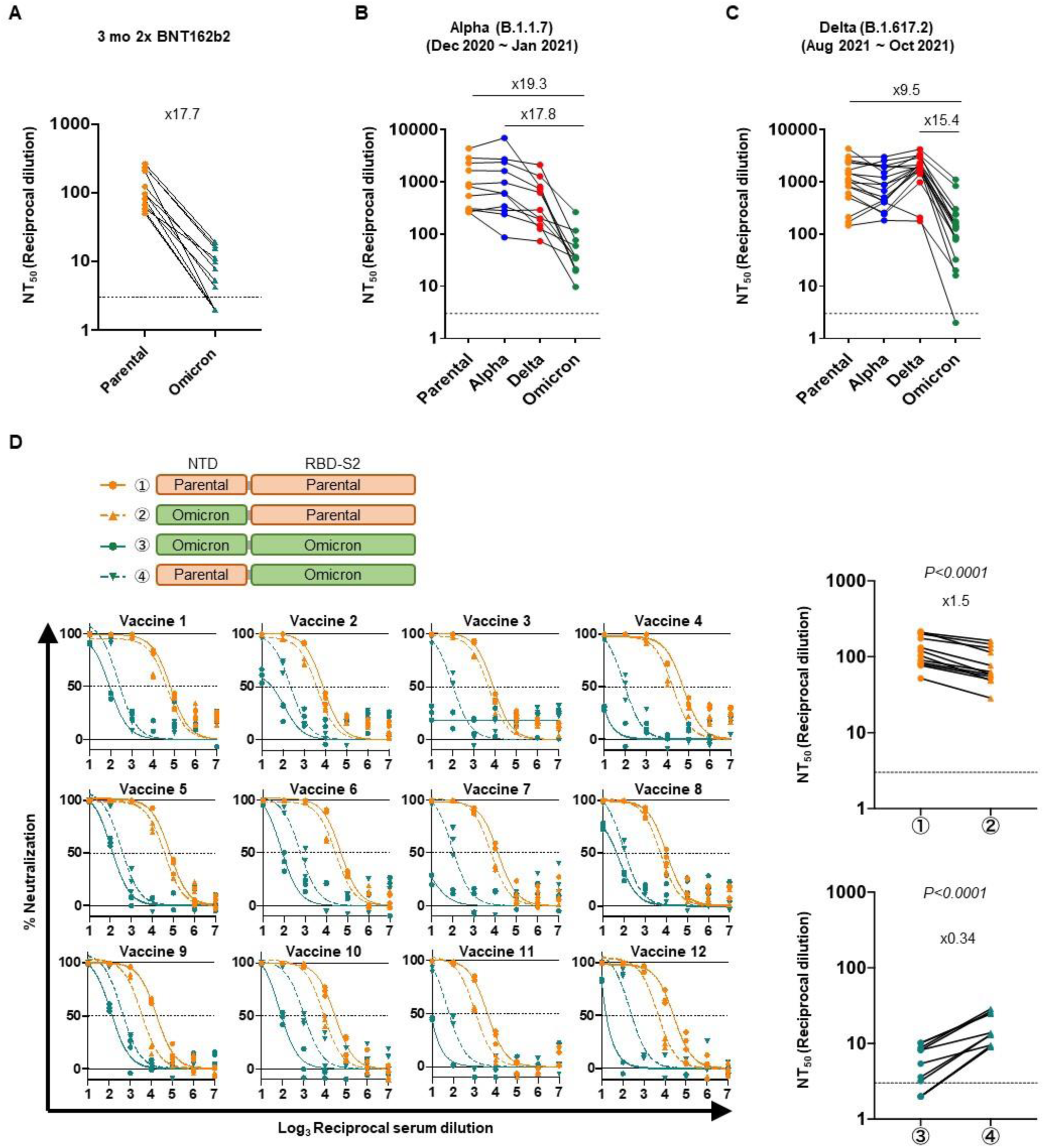
Neutralization assay with Omicron pseudovirus. (**A-C**) The reciprocal dilution of 50% virus neutralization (NT_50_) was determined for sera of 3-month double BNT162b2 vaccination (A), convalescent sera from 11 patients infected with Alpha (B) and 18 patients infected with Delta (C) in 293T/ACE2 cells. Neutralization efficacy of 12 individual vaccinated sera against parental virus (D614G mutation), parental virus with replacement of Omicron’s NTD, Omicron, and Omicron with parental NTD in 293T/ACE2 cells. The NT_50_ was summarized in the right panel. n = 3 technical replicates. *P*-values by two-sided paired *t*-test. Dotted lines indicate detection limit.

To examine the contribution of NTD mutations, we made a pseudovirus harboring a chimeric spike that contained Omicron’s NTD in parental virus or parental NTD in the Omicron variant. Replacing parental virus NTD with Omicron mildly attenuated the neutralizing effect of vaccinated sera. In contrast, the removal of NTD mutations from Omicron generally increased susceptibility to vaccine neutralization (Fig. 2D).

### Engineered ACE2 broadly neutralizes SARS-CoV-2 variants, including Omicron and other sarbecoviruses

We next evaluated the efficacy of neutralizing drugs developed for the original Wuhan strain. Currently a cocktail of imdevimab and casirivimab and a monotherapy of sotrovimab were approved in Japan. The cocktail showed greatly reduced neutralization activity toward Omicron (> 1,000-fold reduction from Wuhan), whereas that of sotrovimab was preserved (Fig. 3A). We previously reported that engineered ACE2 could neutralize a broad range of SARS-CoV-2 variants including Alpha, Beta, and Gamma (*5*). The primary ACE2 mutant in this study, 3N39, carried seven mutations (i.e. A25V, K26E, K31N, E35K, N64I, L79F, and N90H). This mutant was further modified in the later studies to reduce potential immunogenicity, motivating us to remove K26E, N64I, and L79F that made no or little contribution to the enhanced binding, and changed the glycan-eliminating N90H to T92Q, which we reasoned would have a similar effect. Furthermore, the ACE2 collectrin domain (residues 615 to 740) and S128C/V343C were employed to increase productivity and erase enzyme activity, resulting in what we call the 3N39v4 mutant. In the same fashion, 3J320v3 contains the collectrin domain and S128C/V343C. Despite far more extensive mutation in the Omicron RBD compared with these previous VOCs, all three high-affinity engineered ACE2-Fc we developed (3N39v4, 3J113v2, and 3J320v3) exhibited high neutralizing efficacy against Omicron at levels indistinguishable or even higher than the original Wuhan strain (Fig. 3B and Fig. S2). The concept of engineered ACE2 was independently introduced by several groups and their combination of mutations were all unique (*7, 8*). Among these, we tested two different ACE2 mutants reported by Procko and co-workers(*7*), and both showed similar or better neutralization against Omicron (Fig. S2). Wild type ACE2-Fc (740, with the collectrin domain) neutralized Omicron better than Wuhan, similar to Alpha or Delta, which is consistent with a previous report (*9*). Nevertheless, engineered ACE2s maintain an advantage for Omicron over wild-type ACE2 decoy (Fig. 3B).

**Fig. 3.**
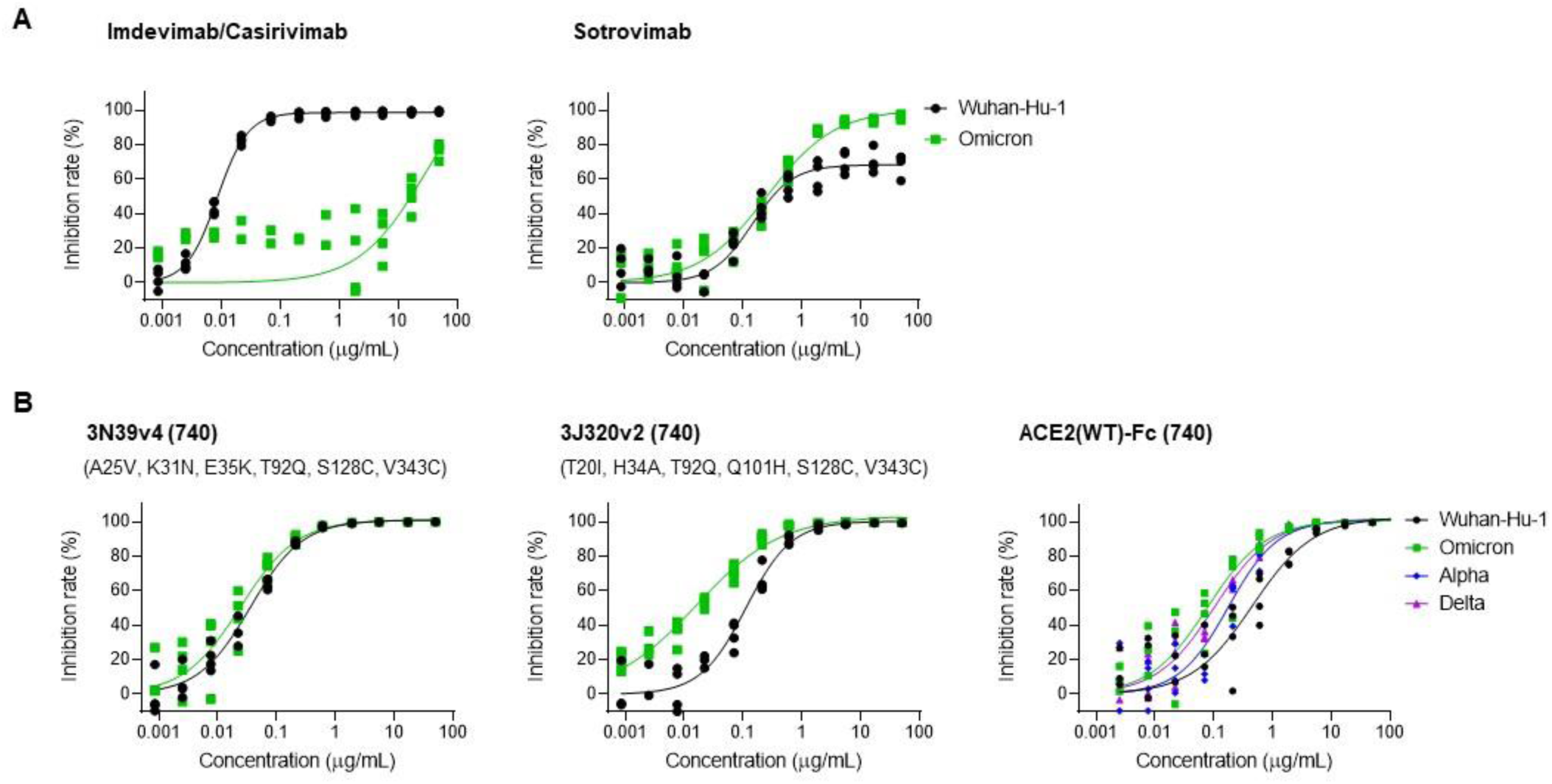
Neutralization assay for monoclonal antibodies and ACE2 decoy. (**A** and **B**) Neutralization efficacy of a cocktail of imdevimab and casirivimab and monotherapy of sotrovimab against Omicron pseudovirus (A) and engineered ACE2 or wild type ACE2 decoy (B) in 293T/ACE2 cells. n = 4 technical replicates. The indicated concentrations of imdevimab and casirivimab were applied in a 1:1 ratio.

We then examined the breadth of cross-neutralization against other SARS-CoV-2 variants and sarbecoviruses by our engineered ACE2 (Fig. 4A and Fig. S3). Previously, we reported the effective neutralization against Alpha, Beta, and Gamma variants as well as SARS-CoV-1(*5*). Here, engineered ACE2 (3N39v4) showed similar or better efficacy to Delta, Epsilon, Lambda or Mu, as compared with the original Wuhan strain (Fig. 4B). Based on the broad representation of the RBD phylogenetic spectrum, 3 different viruses from the SARS-CoV-2 clade and 3 from SARS-CoV-1 clade were also analyzed(*10*) (Fig. 4A and Fig. S3). GD-1, RsSHC014, and WIV1 were tested as pseudoviruses harboring their own spikes, whereas RaTg13, GX-P5L, and Rs4231 were evaluated as chimeric spikes where their RBDs were inserted in the SARS-CoV-1 RBD region in order to obtain the enough infectivity for the analysis (*11*). The high level of inhibition was also observed in GD-1 and WIV1. Engineered ACE2 was less effective in RaTg13, GX, RsSHC014 and Rs4231, but was better than the wild type ACE2 decoy. In contrast, Rs4231 was more sensitive to the wild type decoy (Fig.4C). These results indicate that engineered ACE2 has therapeutic potency against a broad range of SARS-CoV-2 variants and some sarbecoviruses.

**Fig. 4.**
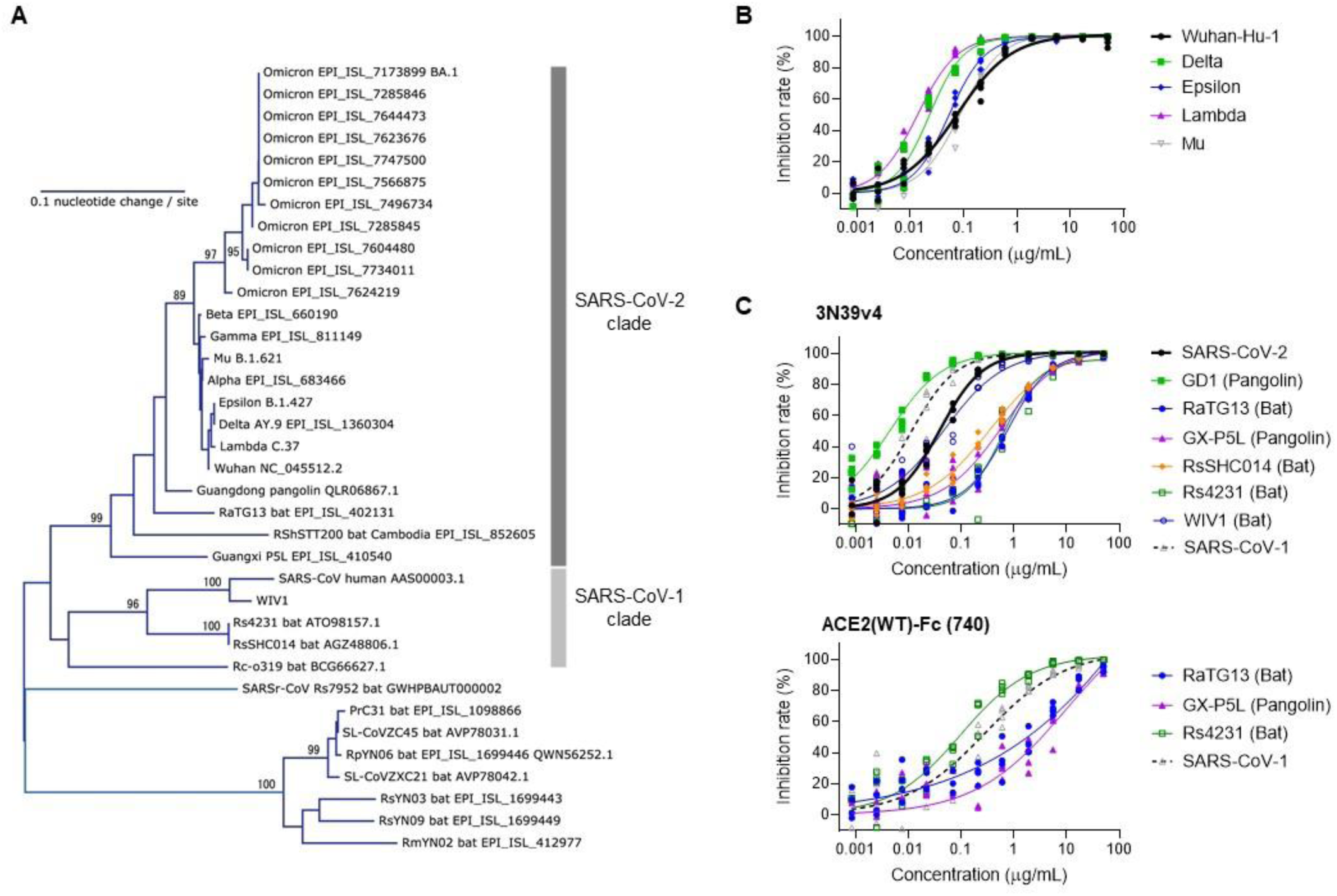
Neutralization assay for engineered with pseudovirus of SARS-CoV-2 variants and sarbecoviruses. (**A**) Phylogenetic tree of the RBD from variants of SARS-CoV-2 and relatives. Accession number or lineage name is shown for each sequence. Bootstrap values larger than 80% are shown at the corresponding branches. (**B**) Neutralization efficacy of engineered ACE2(3N39v4) against previous SARS-CoV-2 variants in 293T/ACE2 cells. (**C**) Neutralization efficacy of engineered ACE2(3N39v4) or wild type ACE2 decoy against sarbecoviruses in 293T/ACE2 cells. n = 4 technical replicates.

### Engineered ACE2 has therapeutic potential against authentic Omicron

Next, we examined the effects of our engineered ACE2 on propagation of authentic Omicron in *in vitro* and *in vivo*. Vero-TMPRSS2 cells were infected with Wuhan or Omicron, together with ACE2-Fc. Consistent with our pseudovirus data, sensitivity of Wuhan or Omicron to engineered ACE2 were comparable (Fig.5A). The therapeutic potential of engineered ACE2 in animal models was reported in both Wuhan and variants (*5, 12*). We thus tested ACE2 against Omicron as well. We infected Syrian hamsters with the virus and treated them with engineered ACE2 2 hours post infection (Fig.5B). After 5 days, viral RNA in the lungs was significantly suppressed by treatment with engineered ACE2 (Fig. 5C). Gene expression of inflammatory cytokines and chemokines showed relative reduction by treatment with engineered ACE2 (Fig. 5D). Omicron-infected hamsters exhibited modest elevation of viral RNA copy number and inflammatory markers, consistent with recent paper reporting mild lung pathology of Omicron infection(*13*). To confirm the efficacy in a more severe model, we examined the effect of engineered ACE2 on survival of CAG-hACE2 mice infected with Omicron. Three out of 4 CAG-hACE2 mice died within 8 days post infection. In contrast, engineered ACE2 treatment significantly rescued these mice (Fig. 5E). These data indicate that our engineered ACE2 retained activity to suppress viral propagation of the Omicron strain.

**Fig. 5.**
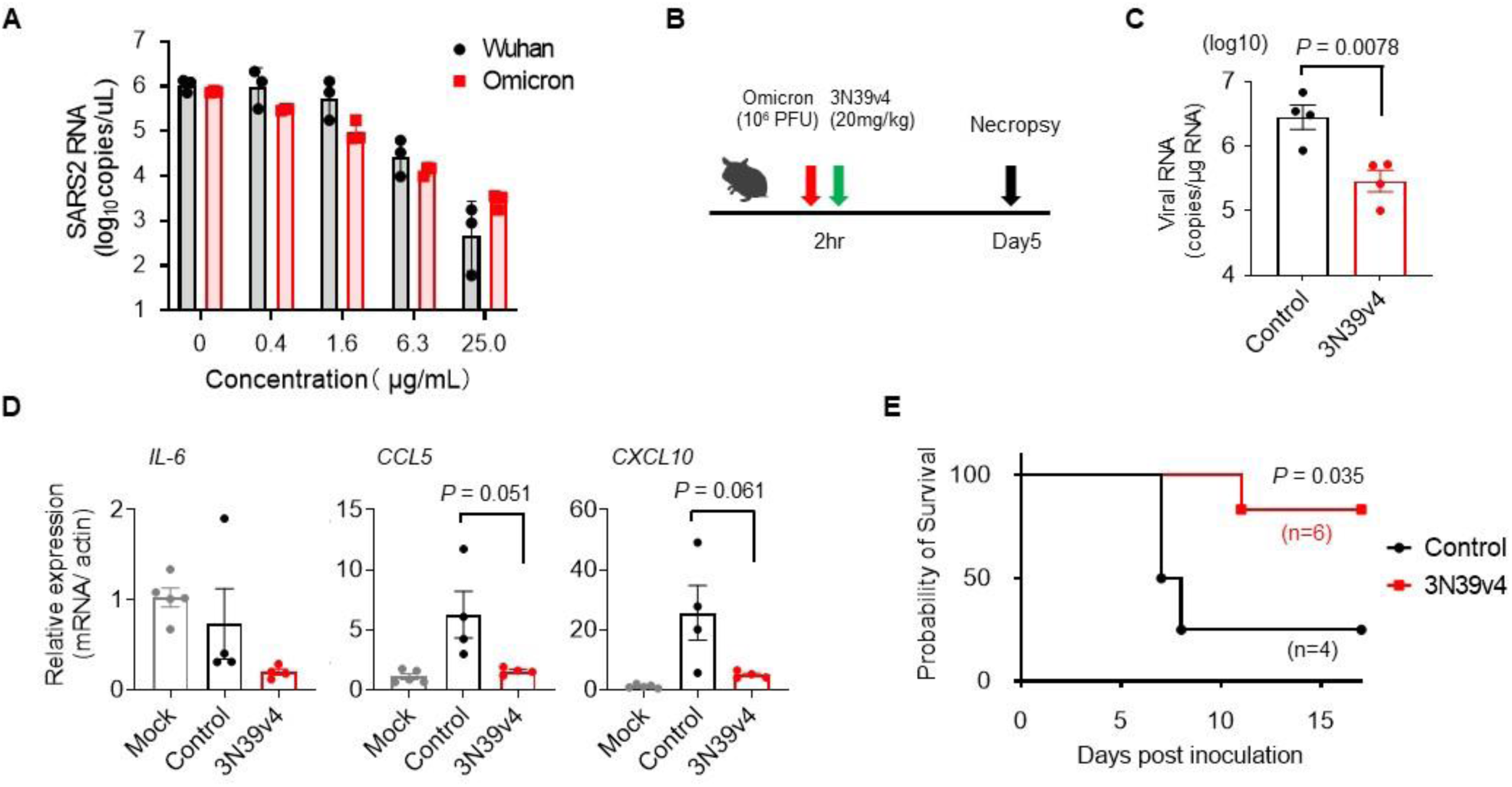
Engineered ACE2 neutralizes authentic Omicron and exhibits therapeutic effect in hamster and hACE2 transgenic mice. (**A**) Sensitivity of engineered ACE2 between the Wuhan and the Omicron strain was compared by using each infectious virus. n = 3 technical replicates. (**B**) Experimental scheme for the hamster model of SARS-CoV-2. (**C**) Quantification of viral RNA in the lungs infected with SARS-CoV-2 Omicron strain was performed by qRT-PCR. (n =4) (**D**) Gene expression of inflammatory cytokines and chemokines was quantified by qRT-PCR. The expression of β-actin was used for normalization. (n =4) The survivals of hACE2-Tg mice infected with the Omicron strain (control: 2 male, 2 female, treated: 3 male and 3 female). *P*-values by two-sided unpaired *t*-test (C and D) and chi-square test (E).

### Structural modeling of the binding mode of engineered ACE2 to Omicron RBD

The structure of the Omicron RBD complexed with wild type ACE2 has recently been determined by several groups (Fig. S4A), revealing sustained binding affinity of this variant compared to the original Wuhan strain via combination of both affinity-enhancing and reducing mutations (*14-16*). We previously determined the crystal structure of Wuhan RBD complexed with the original ACE2 mutant 3N39 (*5*). In the structure, the mutated K35 forms a direct hydrogen bond with Q493 of the Wuhan RBD (Fig. S4B(ii)), whereas the wild type residue E35 forms an intramolecular salt bridge with K31 (Fig.S4B(i), leading us to speculate that the simultaneous mutations of K31N and E35K played a key role in the affinity enhancement of this mutant. On the other hand, Q493 was substituted with Arg in the Omicron RBD, allowing it to make a salt bridge with E35 of the wild type ACE2 (Fig.S4B(iii)). We suspected that this could lower the affinity toward our engineered ACE2 due to electrostatic repulsion with K35. To simulate the binding mode, we built a homology model of a complex between the Omicron RBD (7t9l) and the ACE2 mutant 3N39 (7dmu). In the complex model, the electrostatic clash between K35 of the ACE2 mutant and R493 of Omicron RBD could easily be avoided by side chain rotation. Furthermore, we found that the side chain of R493 could form a direct hydrogen bond with N31 of ACE2(3N39) instead of K35 (Fig. S4B(iv)). Therefore, consistent with the results of the neutralization assay, our engineered ACE2 is expected to have no or little loss of affinity for Omicron RBD.

### No complete escape mutation in deep mutational scanning of the RBD

To comprehensively analyze the alteration of viral infectivity and escape from neutralizing agents, we performed a deep mutational scan (DMS) of the RBD in the context of full-length spike expressed on human Expi293 cells(*17*), followed by incubation with ACE2-harboring GFP-reporter virion. In this “inverted orientation “ setting, ACE2-virion efficiently infected spike-expressing cells only (Fig. S5A). Although DMS analysis on the spike had been done by Bloom and co-workers (*18*), they used yeast surface display of isolated RBDs which would assess the RBD’s structural stability and its affinity against ACE2 or neutralizing agents separately. In contrast, Procko and co-workers used Expi293 cells expressing full-length spike to evaluate the affinity for wild type and engineered ACE2 (*17*). We modified their screening system to monitor the fusion of cell and viral membranes mediated by binding between spike and ACE2, which better mimics the true infection process, albeit in an inverted orientation. The HA-tagged spike library encompassed all 20 amino acid substitutions in the RBD (a.a. P329 to G538) of the original Wuhan spike and was transfected in Expi293 cells in a manner where cells typically acquire no more than a single variant (*17*). After infection of ACE2-harboring virion to library cells with or without neutralizing agents, infected GFP-positive cells and control GFP negative cells were harvested with fluorescence-activated cell sorting (FACS) and the library was sequenced with extracted RNA (Fig.6A). The resulting spike-expressing cells constituted approximately 15% of transfected cells, which is a reasonable rate to avoid multiple library induction (*19*). For infectivity analysis, ACE2-virion was titrated to induce the infection in 2-3% of the cells in consideration of the detection range and library complexity (Fig. S5B). To analyze escape ability, ACE2-virion and neutralization agents were optimized to see the reduction of infectivity from approximately 10% to 3% (Fig. S5B and C). DMS experiments were performed in duplicate, which produced similar results (R^2^ = ∼0.5) as in the case of previous studies conducted on HIV and influenza pathology with library viruses (*20, 21*) (Fig. S6A). We also performed DMS for spike expression level assessed by the staining of HA tagged to N-terminal full-length spike. Among ∼15% HA positive cells, the top 25% and bottom 25% of cells were sorted (Fig. S6B). The resulting single mutant count frequencies correlated well between independent duplicate experiments (R^2^ = 0.93; Fig. S6C). The scan without neutralizing agents provided information about infectivity alteration due to all single amino acid mutations in the RBD (Fig. S7A). When our DMS data for infectivity and expression were compared with those obtained in the previous yeast surface display DMS from the Bloom group (*18*), infectivity was mildly correlated with Bloom’s affinity data that was based on ACE2-binding in several ACE2 concentrations (R^2^ = 0.23). In contrast, expression data correlated better, in spite of the difference in using full-length spike and isolated RBD (R^2^ = 0.48; Fig. S7B-E).

According to the scanning data for infectivity, 8 of 15 mutations in the RBD of Omicron may enhance infectivity, whereas 3 mutations (S371L, S373P, S375F) that exist in the conserved region may reduce infection (Fig.6B). Incubation of library cells and ACE2 virion with neutralizing drug and sera succeeded in revealing the mutational patterns that enable evasion. For example, the scan in the presence of casirivimab detected all epitope residues previously reported (*22, 23*), which confirmed that this assay had adequate sensitivity to detect actual escape mutations (Fig.6C and Fig. S8B). In this panel, Omicron mutation, K417N, E484A and Q493R were all indicated to contribute to escape from casirivimab (Fig.6C). Some VOCs exhibited partial resistance to neutralization by immunized sera(*24*); however, actual escape is achieved by a combination of mutations, not by any single mutation (*25*). Consistently, the scan measuring resistance to vaccinated sera revealed that alteration of escape value simply paralleled that of infectivity without irregular enhancements, unlike the case of casirivimab (Fig.6D and Fig. S8C). This result was reproduced by the sera from a different vaccinated individual (Fig. S9). A similar result was also observed in a scan for escape from engineered ACE2 (Fig. 6E and Fig. S8D). When major escape value mutants obtained through the DMS were individually subjected to the neutralization assay, it was found that L455Y and N487Q partially reduced the neutralization efficacy in both 3N39v4 and wild type ACE2 decoys (Fig. 6F). However, engineered ACE2 still maintained a neutralization efficacy similar to the level of sotrovimab for Wuhan and Omicron (Fig. 3A). These results provide evidence that engineered ACE2 neutralizes each single-residue mutation more effectively than monoclonal antibodies.

**Fig. 6.**
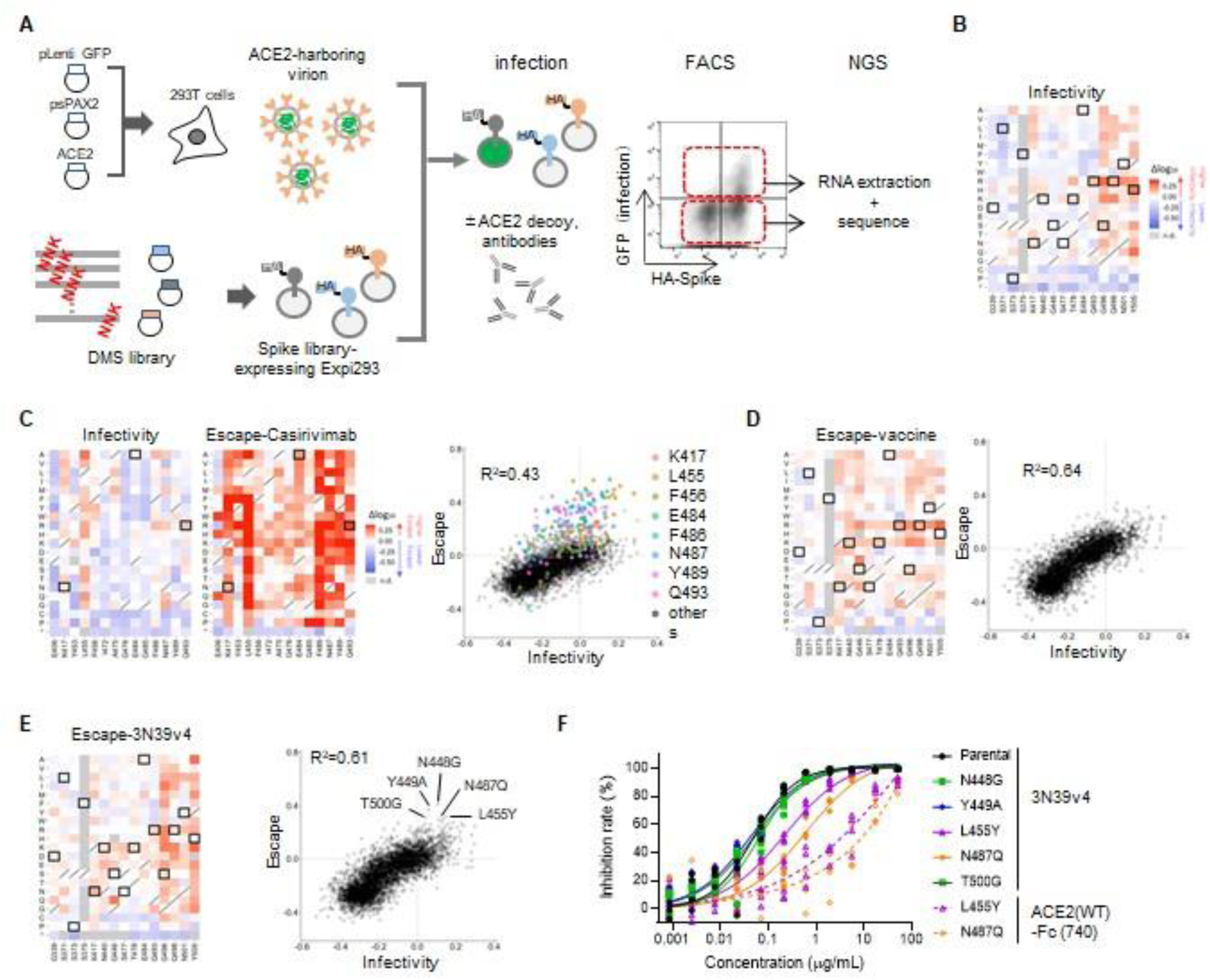
Deep mutational scan identified no single residue mutation to induce escape from engineered ACE2. (**A**) Schematic of the deep mutational scanning to evaluate infectivity and escape from neutralizing agents. (**B**) Heatmaps illustrating how all single mutations that Omicron obtains affect its infectivity. Open squares are Omicron-carrying mutation. (**C**) Heatmaps showing the alteration of infectivity and escape value from casirivimab in casirivimab antigen sites of the spike protein. Open squares are Omicron-carrying mutation (left). Correlation in mutation effects on infectivity and escape from casirivimab. Color dots are all amino acid substitution at the indicated major casirivimab antigen sites (right). (**D**) Heatmaps illustrating how all single mutations that Omicron obtains affect its escape from vaccinated sera Open squares are Omicron-carrying mutation (left). Correlation in mutation effects on infectivity and escape from vaccinated sera (right). (**E**) Heatmaps illustrating how all single mutations that Omicron obtains affect its escape from 3N39v4. Open squares are Omicron-carrying mutation (left). Dot plots showing the correlation between infectivity and escape value from 3N39v4. Indicated mutants were individually analyzed in the next. (**F**) Neutralization efficacy of 3N39v4 or wild type ACE2 decoy against 5 escape candidates in 293T/ACE2 cells. n = 4 technical replicates. Heatmap squares are colored by mutational effect according to scale bars on the right, with blue indicating deleterious mutations. Heatmaps of the whole spike are available in supplementary figure 7 and 8.

## DISCUSSION

Omicron contains approximately 30 mutations in the spike protein and escapes vaccine, convalescent sera, and some therapeutic antibodies (*1-4, 26-29*). Intensive analyses from all over the world indicate approximately a 10 to 50-fold reduction in the neutralization titer for Omicron in vaccines and convalescent subjects from various variants (*26, 27, 29*). Consistently, our assay using Omicron pseudovirus revealed that 17.7-fold and 9.5 to 19.3-fold reduction in sera from 2x BNT162b2 vaccinated and previously infected people, respectively. The assay using pseudovirus harboring chimeric spike of Omicron and the original Wuhan strain intriguingly demonstrated that multiple mutations in the NTD could contribute to reduce neutralization in addition to major escape due to RBD mutations. Previous studies showed that the deletion mutation in the NTD induced escape from isolated NTD-binding neutralizing antibodies (*30*). According to another report, vaccinated and convalescent sera exhibited weak neutralization titers, even after depletion of RBD-binding antibodies (*31*). Furthermore, direct structural analysis of NTD-targeted neutralizing antibodies indicated that they each recognize a common glycan-free, electropositive surface comprised of flexible loops where most Omicron NTD mutations are located (*6*). These reports point to the possibility of NTD-mediated active neutralization by immunized sera. Consistent with this idea, the present study demonstrates that the extensive escape of Omicron can be achieved, in part, by attenuated neutralization of NTD-binding antibodies. Regarding the escape due to RBD mutations, immunized sera contain polyclonal antibodies and are basically less affected by single mutations. However, a previous DMS study identified RBD positions F456 and E484 mutated to alanine in Omicron, as major epitopes of immunized sera (*31*).

For therapeutic strategies to overcome virus mutation, some antibodies are used in the form of a cocktail and others are designed to target a conserved region, in order to prevent mutational escape. The present study showed that a cocktail of imdevimab and casirivimab failed to neutralize Omicron, whereas sotrovimab, which targets a highly conserved epitope, remained effective. Other papers also reported impaired neutralization in a wide range of monoclonal antibodies (*2-4, 28*), and that only sotrovimab was minimally affected among those in clinical use (*2*). Omicron exhibits the ability to evade from combinations of antibodies and acquired mutations even in highly conserved sites (S371L, S373P, S375F). These observations provide a sobering outlook for the therapeutic development of monoclonal antibodies against viruses in the context of escape mutation. In contrast, engineered ACE2 successfully neutralized Omicron and even other sarbecoviruses that could form the basis of future SARS-CoV-3. The emergence of Omicron reinforces the difficulty of drug development against viral infection and highlights the strength of receptor decoys as a therapeutic strategy to counteract COVID-19 mutational escape.

In this study, we performed DMS using full-length spike-expressing human cells and ACE2-harboring virion. Incubation of these components aimed to functionally reproduce actual infection of host cells. In contrast, Bloom performed DMS with yeast surface display and analyzed RBD expression levels and binding of RBD with several concentrations of ACE2. Yeast screening has the advantage in the library size and the ability to restrict incorporation of multiple mutants due to the exclusive nature of different plasmids. Mammalian cell-based screening has a limit in library size and can suffer from contamination of multiple mutant expression; however, its strength is that it allows analysis of various phenotypes, including virus infection. When compared with Bloom’s DMS, the pattern of effects on protein expression levels showed similar trends, in spite of difference in the form of spike used (isolated RBD and full-length spike). On the other hand, our infectivity DMS was less correlated with Bloom’s ACE2 affinity data. One clear explanation for this difference was the assayed phenotypes themselves; however, further discussion requires intensive validation of each DMS system (Fig. S7E). Through DMS, we found that L455Y and N487Q mildly reduced neutralization efficacy of engineered ACE2 (Fig. 6F). These residues make direct contacts with ACE2 but are away from the affinity-enhancing mutations of engineered ACE2 (*5*). As a result, these mutations also caused the reduction of the neutralization ability of wild type ACE2 in a similar manner. In the case of N487Q, both DMS results showed elevated infectivity, increased expression levels, and reduced ACE2-binding affinity. It is thus possible that its reduced ACE2 affinity is offset by the increased spike expression/stability to maintain the overall infectivity, while causing attenuation of the neutralization ability of ACE2 decoys. Another study also reported that certain RBD mutations affected other high-affinity ACE2 decoys (*32*). In theory, similar viral mutations that reduce the neutralization efficacy of engineered ACE2 may arise. However, engineered ACE2 decoys were not obtained through natural evolution in the presence of an actual immune system; indeed, natural virus would not encounter such agents unless they were actually used as a therapeutic intervention. In contrast, viruses circulating in the wild are under selection pressure to escape antibody-based therapeutics due to vaccination and prior infections. Therefore, preferential emergence of ACE2 escape mutants outside the lab is expected to be rare, making the ACE2 decoy a highly valid therapeutic modality against this recalcitrant virus.

## MATERIALS AND METHODS

### Ethics statement

Blood samples were obtained from hospitalized adults with PCR-confirmed SARS-CoV-2 infection and vaccinated individuals who were enrolled in a prospective cohort study approved by the Clinical Research Review Committee in Kyoto Prefectural University of medicine (ERB-C-1810-2, ERB-C-1949-1). All human subjects provided written informed consent.

All animal experiments with SARS-CoV-2 were performed in biosafety level 3 (ABSL3) facilities at the Research Institute for Microbial Diseases, Osaka University. Animal Experimentation, and the study protocol was approved by the Institutional Committee of Laboratory Animal Experimentation of the Research Institute for Microbial Diseases, Osaka University (R02-08-0). All efforts were made during the study to minimize animal suffering and to reduce the number of animals used in the experiments.

### Human sera

Peripheral blood was collected from 12 persons (7 males, 5 females, age 24-56 mean 37.5) 3 months after vaccination with double Pfizer-BioNTech vaccine BNT162b2, 11 convalescents (7 males, 4 females, age 50-83 mean 69) before the Delta pandemic (December 2020 through January 2021), and 18 convalescents (15 males, 3 females, age 36-77 mean 56.6) during/after the Delta pandemic (August 2021 through October 2021). The sera were isolated and stored at –80°C until use.

### Omicron genetic and epidemic analysis

Amino acid mutation frequencies in the spike protein for each variant (Alpha = 1,138,704, Beta = 40,135, Gamma = 117,200, Delta = 3,441,137, and Omicron = 5,469 sequences) were extracted from the report of outbreak.info (https://outbreak.info/compare-lineages?pango=Delta&pango=Omicron&pango=Alpha&pango=Beta&pango=Gamma&gene=S&threshold=0.2&nthresh=1&sub=false&dark=true%29.%20Accessed%2017%20December%202021.) as of December 17, 2021. Mutation frequencies of each amino acid substitution which was observed in Omicron were plotted using “corrplot ” package in R. Time course of variant distribution was analyzed by Nextclade ver 1.7.0 (https://joss.theoj.org/papers/10.21105/joss.03773) from SARS-CoV-2 nucleic acid sequences which were downloaded from GISAID database as of December 17, 2021. A phylogenetic tree of spike protein from SARS-CoV-2 variants and relatives were inferred based on full-length amino acid sequences taken from GISAID, genbank, National Genomics Data Center and the database of SARS-CoV-2 variants by Stanford HIVDB team. The neighbor joining method(*33*) was applied to a distance matrix estimated by the maximum likelihood method. To perform multiple sequence alignment of spike proteins 6,000,693 spike protein amino acid sequences were downloaded from GISAID on Dec 14, 2021. The frequency of all unique RBD regions (defined as R319-541F) were determined for each major variant (as defined in https://en.wikipedia.org/wiki/Variants_of_SARS-CoV-2), and the most frequent instance within a variant was used as a representative. The sequences were multiply aligned by MAFFT(*34*) with the corresponding RBD regions from PG-GD1, VIW1 and CoV-1. The MSA was rendered using an in-house script to show only positions non-identical to the Wuhan strain.

### Cell culture

Lenti-X 293T cells (Clontech) and its derivative, 293T/ACE2 cells were cultured at 37 °C with 5% CO_2_ in Dulbecco’s modified Eagle’s medium (DMEM, WAKO) containing 10% fetal bovine serum (Gibco) and penicillin/streptomycin (100 U/ml, Invitrogen). VeroE6/TMPRSS2 cells were a gift from National Institutes of Biomedical Innovation, Health and Nutrition (Japan) and cultured at 37 °C with 5% CO2 in DMEM (WAKO) containing 5% fetal bovine serum (Gibco) and penicillin/streptomycin (100 U/ml, Invitrogen). All the cell lines were routinely tested negative for mycoplasma contamination.

### Protein synthesis and purification

Monoclonal antibodies and engineered ACE2 were expressed using the Expi293F cell expression system (Thermo Fisher Scientific) according to the manufacturer’s protocol. Fc-fused proteins were purified from conditioned media using the rProtein A Sepharose Fast Flow (Cytiva), respectively. Fractions containing target proteins were pooled and dialyzed against phosphate buffered saline (PBS).

### Pseudotyped virus neutralization assay

Pseudotyped reporter virus assays were conducted as previously described (*5*). With a plasmid coding SARS-CoV-2 Spike (addgene #145032) as a template, Omicron and other variants mutations and ΔC19 deletion (with 19 amino acids deleted from the C terminus) was cloned into pcDNA4TO (Invitrogen) (*35*). Pangolin CoV GD-1, Bat CoV RsSHC014, and WIV1 spikes were synthesized (Integrated DNA Technologies) and cloned into pcDNA4TO (Invitrogen) in the form of ΔC19(*10, 36*). Pangolin CoV GX-P5L, Bat CoV RaTG13, and Rs4231 RBDs were synthesized (Integrated DNA Technologies) and cloned into the RBD of SARS-CoV-1 (ΔC19) (*5, 10, 11, 36*). Spike-pseudovirus with a luciferase reporter gene was prepared by transfecting plasmids (OmicronΔC19, psPAX2-IN/HiBiT (*37*), and pLenti firefly) into LentiX-293T cells with Lipofectamine 3000 (Invitrogen). After 48 hr, supernatants were harvested, filtered with a 0.45 μm low protein-binding filter (SFCA), and frozen at –80 °C. The 293T/ACE2 cells were seeded at 10,000 cells per well in 96-well plate. HiBit value-matched Pseudovirus and three-fold dilution series of serum or therapeutic agents were incubated for 1 hr, then this mixture was administered to ACE2/293T cells. After 1 hr pre-incubation, medium was changed and cellular expression of luciferase reporter indicating viral infection was determined using ONE-Glo™ Luciferase Assay System (Promega) in 48 hr after infection. Luminescence was read on Infinite F200 pro system (Tecan). The assay of each serum was performed in triplicate, and the 50% neutralization titer was calculated using Prism 9 (GraphPad Software).

### Library construction, FACS, and Illumina sequencing analysis

Saturation mutagenesis was focused on the original Wuhan strain spike residues F329 to C538 forming the RBD. Pooled oligo with degenerate NNK codon was synthesized by Integrated DNA Technologies, Inc. Synthesized oligo was extended by overlap PCR and cloned into pcDNA4TO HMM38-HA-full length S plasmid. Transient transfection conditions were used that typically provide no more than a single coding variant per cell(*17*). Expi293F cells at 2 × 10^6^/ml were transfected with a mixture of 1 ng of library plasmid with 1 μg of pMSCV as a junk plasmid per ml using ExpiFectamine (Thermo Fisher Scientific). Twenty-four hours after transfection, cells were incubated with ACE2-harboring GFP reporter virion. In case of escape analysis, cells were pre-incubated with neutralizing agents for 1 hr. One-hour incubation with ACE2 virion, the medium was replaced and cells were collected 24 hours after infection for FACS. Cells were washed twice with PBS-BSA and then co-stained for 20 min with anti-HA Alexa Fluor 647 (clone TANA2,1/4000 dilution; MBL). Cells were again washed twice before sorting on a MA900 (Sony). Dead cells, doublets, and debris were excluded by first gating on the main population by forward/side scattering. From the HA positive (Alexa Fluor 647) population, GFP-positive and -negative cells were collected. The total numbers of collected cells were ∼2 million cells for each group. Total RNA was extracted from collected cells TRIzol (Life Technologies) and Direct-zol RNA MiniPrep (Zymo Research Corporation) according to the manufacturer’s protocol. First-strand complementary DNA (cDNA) was synthesized with Accuscript (Agilent) primed with a gene-specific oligonucleotide. Library was designed for 3 sections separately in the RBD and pooled. After cDNA synthesis, each library was amplified with specific primers. Following a second round of PCR, primers added adapters for annealing to the Illumina flow cell and sequencing primers, together with barcodes for experiment identification. The PCR products were sequenced on an Illumina NovaSeq 6000 using a 2 × 150 nucleotide paired-end protocol in Department of Infection Metagenomics, Research Institute for Microbial Diseases, Osaka University. Data were analyzed comparing the read counts with each group normalized by wild type sequence read-count. Log10 enrichment ratios for all the individual mutations were calculated and normalize by subtracting the log10 enrichment ratio for the wild type sequence across the same PCR-amplified fragment.

### Viruses

SARS-CoV-2 (Wuhan: 2019-nCoV/Japan/TY/WK-521/2020, Omicron: 2019-nCoV/Japan/TY38-873/2021) strain was isolated at National Institute of Infectious Diseases (NIID). SARS-CoV-2 were propagated in VeroE6-TMPRSS2 cells. Viral supernatant was harvested at two days post infection and the viral titer was determined by plaque assay.

### SARS-CoV-2 neutralization assay

Vero-TMPRSS2 were seeded at 80,000 cells in 24 well plates and incubated for overnight. Then, SARS-CoV-2 was infected at MOI of 0.1 together with sACE2-Fc protein. After 2 hr, cells were washed by fresh medium and incubated with fresh medium for 22 hr. Culture supernatants were collected and performed qRT-PCR assay.

### Animal models of SARS-CoV-2 infection

Four weeks-old male Syrian hamsters were purchased from SLC Japan. Syrian hamsters were anaesthetized by intraperitoneal administration of 0.75 mg kg-1 medetomidine (Meiji Seika), 2 mg kg-1 midazolam (Sandoz) and 2.5 mg kg-1 butorphanol tartrate (Meiji Seika) and challenged with 1.0 × 10^4^ PFU (in 60μL) via intranasal routes. After 2 hr post infection, ACE2-Fc (3N39v2, 20mg kg-1) were dosed through intraperitoneal injection. On 5 days post infection, all animals were euthanized and lungs were collected for histopathological examinations, virus titration and qRT-PCR.

CAG-hACE2 transgenic mice (hACE2-Tg) were obtained from the Laboratory Animal Resource Bank of the National Institutes of Biomedical Innovation, Health and Nutrition (NIBIOHN). These mice were maintained by crossing wild type C57BL/6 mice. The primers of genotyping were 5 ‘-CTTGGTGATATGTGGGGTAGA-3 ‘ and 5′-CGCTTCATCTCCCACCACTT-3′ as shown previously (PMID:

34463644). Male and female mice aged at four-week-old were anaesthetized and challenged with 1.0 × 10^3^ PFU (in 20μL) via intranasal routes. After 2 hr post infection, ACE2-Fc (3N39v2, 20mg kg-1) were dosed through intravenous injection and monitored their survivals.

### Quantitative RT-PCR

Total RNA of lung homogenates was isolated using ISOGENE II (NIPPON GENE). Real-time RT-PCR was performed with the Power SYBR Green RNA-to-CT 1-Step Kit (Applied Biosystems) using a AriaMx Real-Time PCR system (Agilent). The relative quantitation of target mRNA levels was performed by using the 2-ΔΔCT method. The values were normalized by those of the housekeeping gene, β-actin. The following primers were used: for β-actin; 5 ‘-TTGCTGACAGGATGCAGAAG-3 ‘ and 5 ‘-GTACTTGCGCTCAGGAGGAG-3 ‘, 2019-nCoV_N2; 5 ‘-AAATTTTGGGGACCAGGAAC -3 ‘and 5 ‘-TGGCAGCTGTGTAGGTCAAC -3 ‘, IL-6; 5 ‘-GGA CAATGACTATGTGTTGTTAGAA -3 ‘and 5 ‘-AGGCAAATTTCCCAATTGTATCCAG -3 ‘, CCL5; 5 ‘-TCAGCTTGGTTTGGGAGCAA -3 ‘and 5 ‘-TGAAGTGCTGGTTTCTTGGGT -3 ‘, CXCL10; 5 ‘-TACGTCGGCCTATGGCTACT -3 ‘and 5 ‘-TTGGGGACTCTTGTCACTGG -3 ‘.

### Quantitative RT-PCR of Viral RNA in the supernatant

The amount of RNA copies in the culture medium was determined using a qRT-PCR assay as previously described with slight modifications (*38*). In brief, 5 μl of culture supernatants were mixed with 5 μl of 2 × RNA lysis buffer (2% Triton X-100, 50 mM KCl, 100 mM Tris-HCl [pH 7.4], 40% glycerol, 0.4 U/μl of Superase·IN [Life Technologies]) and incubate at room temperature for 10 min, followed by addition of 90 μl of RNase free water. 2.5 μl of volume of the diluted samples was added to 17.5 μl of a reaction mixture. Real-time RT-PCR was performed with the Power SYBR Green RNA-to-CT 1-Step Kit (Applied Biosystems) using a AriaMx Real-Time PCR system (Agilent).

### Statistical analysis

Neutralization measurements were done in technical triplicates and relative luciferase units were converted to percent neutralization and plotted with a non-linear regression model to determine IC50 values using GraphPad PRISM software (version 9.0.0). Comparisons between two groups were made with paired *t* test, unpaired *t* test, and chi-square test.

## SUPPLEMENTARY MATERIALS

Figs. S1 to S9

## ACKNOWLEDGEMENTS

We would like to thank Takaaki Nakaya (Kyoto Prefectural University of Medicine) for helpful discussion; Kenzo Tokunaga (Department of Pathology, National Institute of Infectious Diseases) for kind gift of plasmid coding psPAX2-IN/HiBiT, Kiyoshi Tanabayashi (Department of Veterinary Science, National Institute of Infectious Diseases) for the use of CAG-hACE2 transgenic mice. Funding: This work was supported by Japan Agency for Medical Research and Development (AMED), Research Program on Emerging and Re-emerging Infectious Diseases under JP 21fk0108465h0001 (A.H., J.T. and T.O.), Platform Project for Supporting Drug Discovery and Life Science Research (Basis for Supporting Innovative Drug Discovery and Life Science Research) under JP21am0101075 (J.T.), JP21am0101108 (D.M.S) and grant from Mochida Memorial Foundation for Medical and Pharmaceutical Research (A.H), SENSHIN Medical Research Foundation (N.I).

## Author contribution

A.H. designed the research; N.I. and Y.H. performed pseudovirus neutralization assay; N.I, S.T. and Y.H. performed deep mutational analysis; T.I. provided clinical samples of vaccinated and convalescent sera; Y.K., D.M., Y.O., and S.N. performed and analyzed next-generation sequencing; T.A., J.T. purified and prepared the proteins; T.A. performed structure analysis; K.K., S.L. and D.M.S. conducted phylogenetic analysis and manage SpikeAtlas database; Y.I., T.S. and T.O. conducted authentic virus experiments; D.M.S., T.O., J.T., and A.H. supervised the research; A.H., J.T., and T.O. wrote the manuscript; all authors discussed the results and commented on the manuscript.

## Data and materials availability

All unique/stable reagents generated in this study are available with a completed Materials Transfer Agreement. Deep mutational scan data has been deposited at SpikeDB (https://sysimm.ifrec.osaka-u.ac.jp/sarscov2_dms/). Structural data of S trimer and ACE2 complex, wild type ACE2/Wuhan-RBD complex, wild type ACE2/Omicron-RBD complex and ACE2 (3N39)/Wuhan-RBD complex are derived from PBD codes: 7A89, 6M0J, 7T9L, and 7DMU, respectively. DMS data for affinity against ACE2 and expression are retrieved from GitHub (https://github.com/jbloomlab/SARS-CoV-2-RBD_DMS).

## Supplementary Materials

**Fig. S1.**
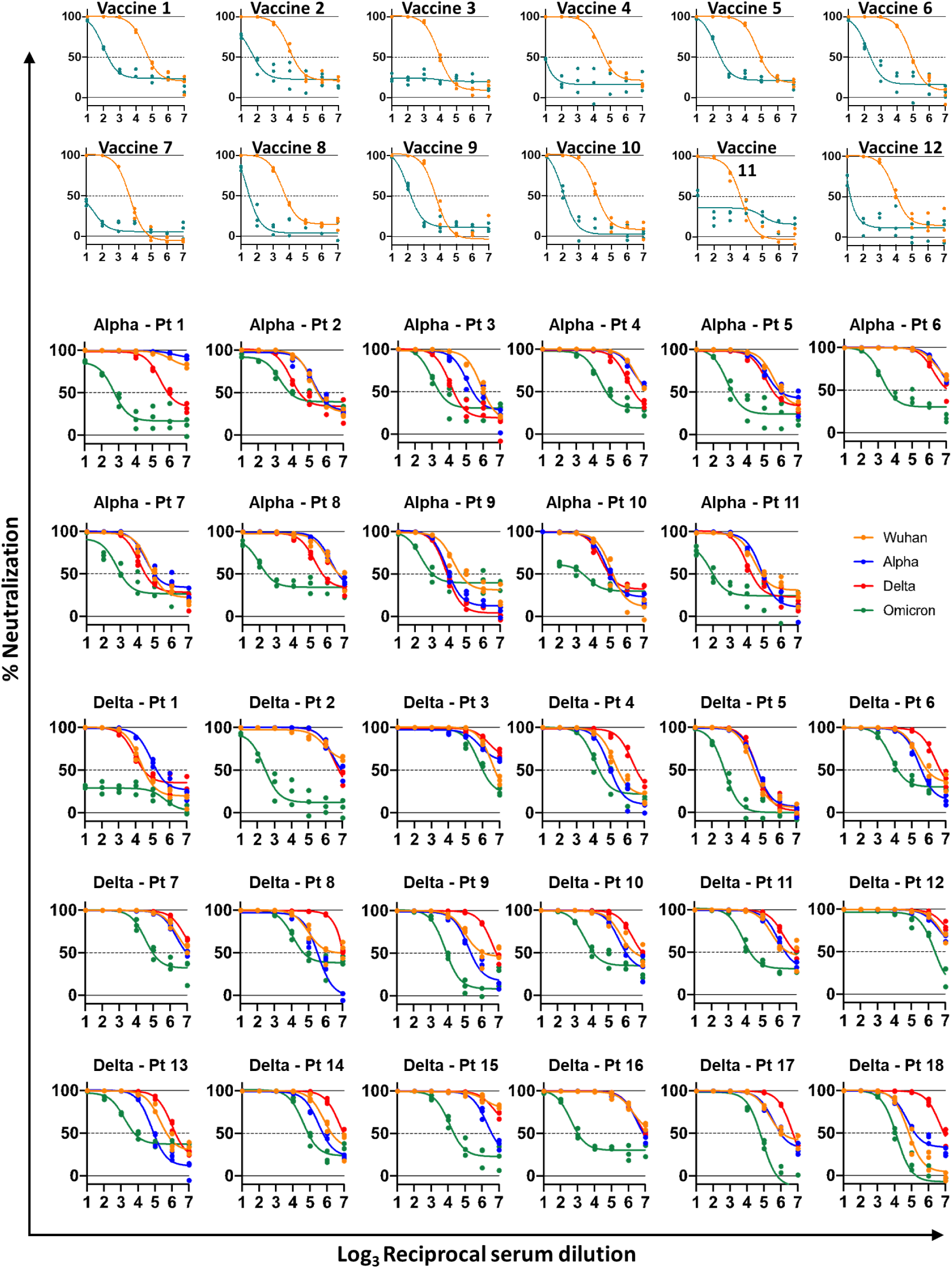
Neutralization assay with Omicron pseudovirus. Individual data for Figure 1A to 1C. Neutralization of the parental (D614G) pseudovirus (red), Alpha (blue), Delta (red) and Omicron (green) analyzed in 293T/ACE2 cells. n = 3 technical replicates.

**Fig. S2.**
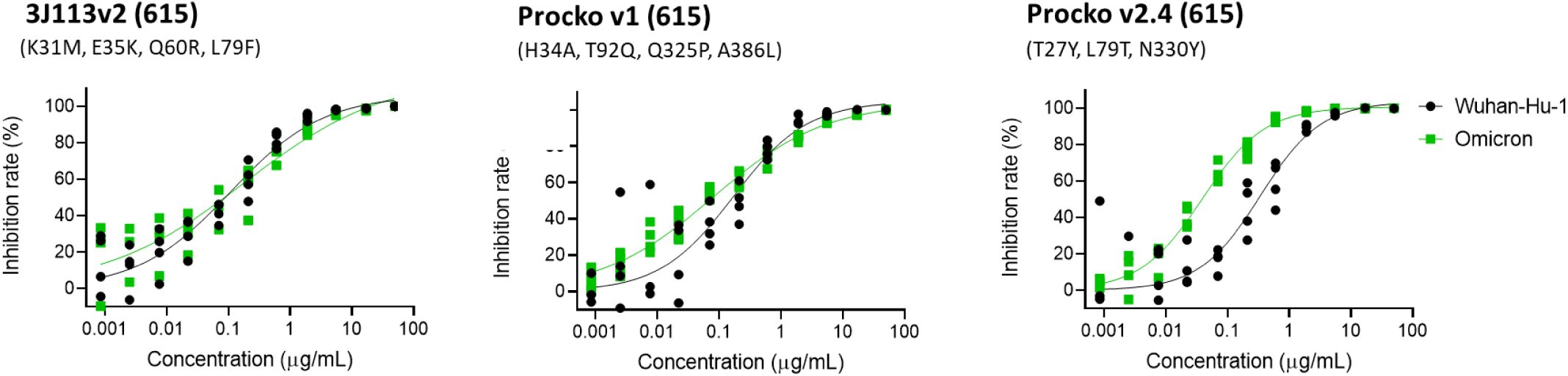
Neutralization assay for engineered ACE2s. Neutralization efficacy of 3J113v2 from our previous study and v1 and v2.4 in the paper reported by Procko (ref 7); three different ACE2-Fc (615 amino acid, without the collectrin domain) against Omicron pseudovirus analyzed in 293T/ACE2 cells. n = 4 technical replicates.

**Fig. S3.**
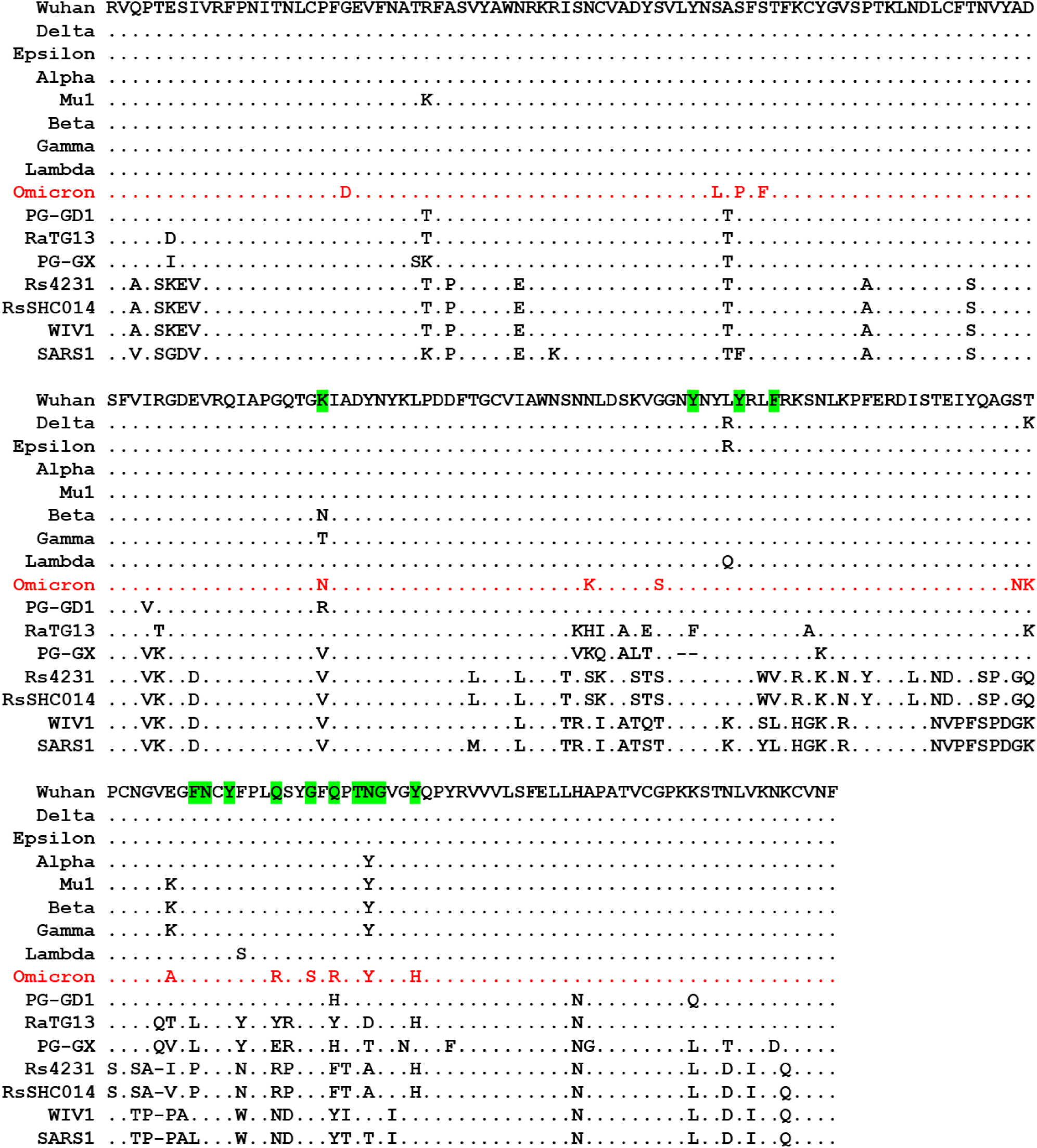
Alignment of amino acid sequences of sarbecovirus RBDs. ACE2 contacts are highlighted by green based on the crystal structure (PDB: 6M0J).

**Fig. S4.**
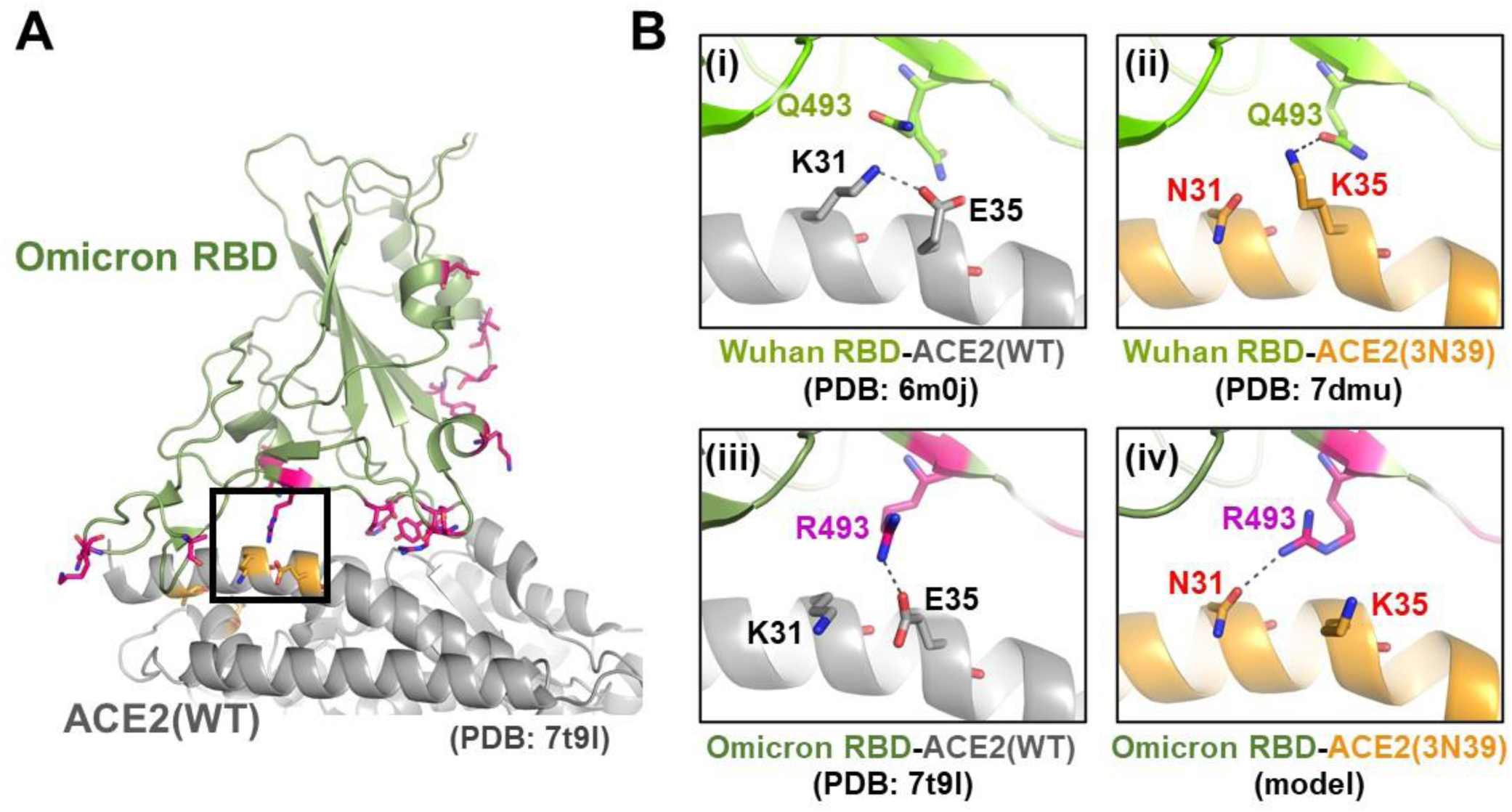
Difference in the interaction mode of Wuhan RBD and Omicron RBD to ACE2 mutant. (**A**) Cryo-electron microscopy structure of the wild type ACE2-Omicron RBD complex (Manner et al, PDB ID: 7t9i). Residues mutated in Omicron are shown as magenta stick models. The ACE2 residues mutated in our mutant 3N39v4 (only A25, K31, E35, and T92 are close to the interface and visible in this view) are shown in pale orange. **(B)** Close-up views of the interface between (i) Wuhan RBD-WT ACE2 (PDB ID: 6m0j), (ii) Wuhan RBD-3N39 ACE2 (PDB ID: 7dmu), (iii) Omicron RBD-WT ACE2 (PDB ID: 7t9l), and (iv) Omicron RBD-3N39 ACE2 (simulated model). Potential hydrogen-bonding/salt-bridge interactions are indicated by dashed lines.

**Fig. S5.**
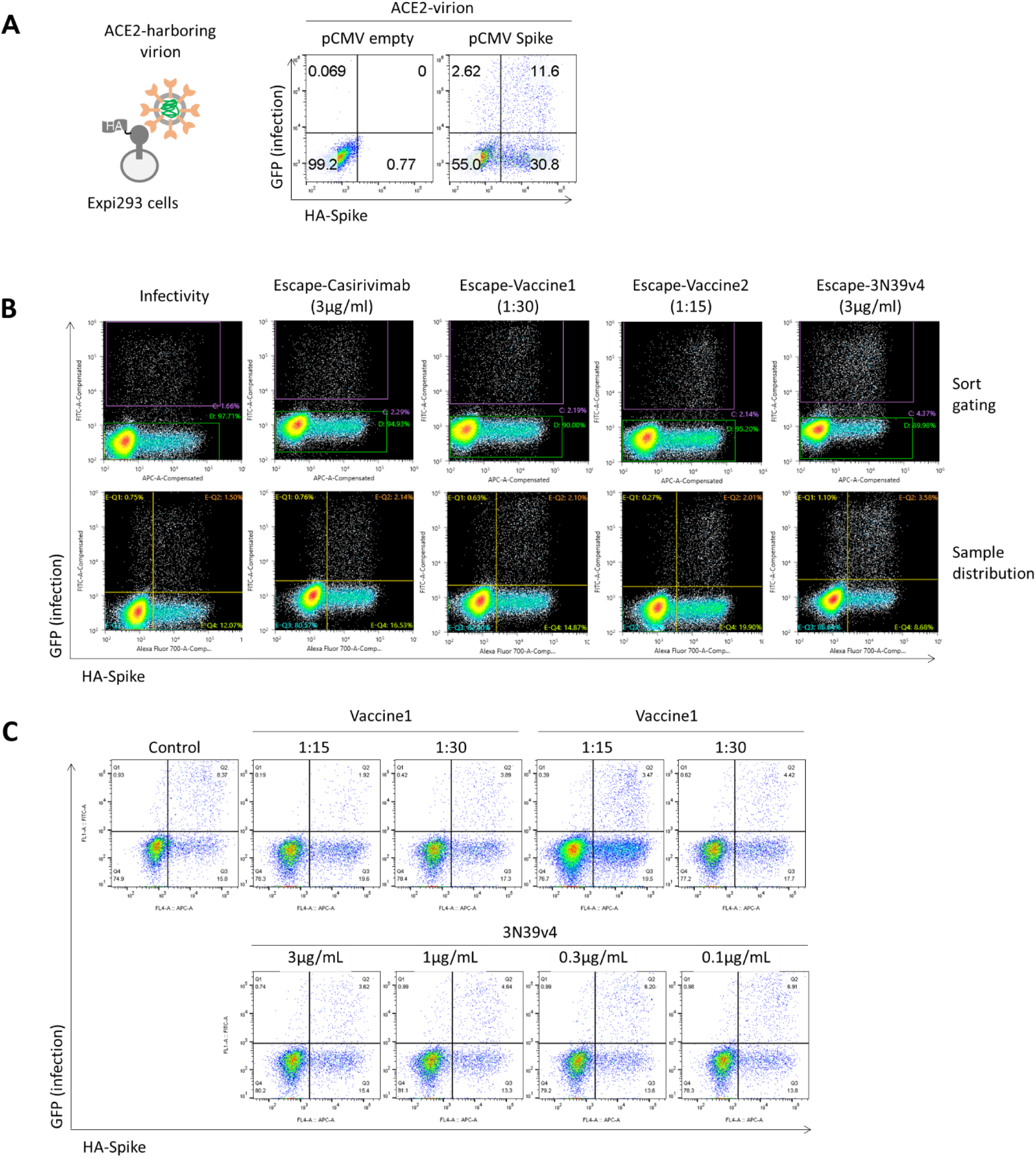
DMS for infectivity and escape using spike-expressing cells and ACE2-harboring virion. (**A**) Flowcytometry showing that ACE2-virion infected only spike expressing cells. **(B)** Upper panels are the gate setting to sort infected cells (gate C) and non-infected cells (gate D). Lower panels show the proportion of each population. **(C)** The titration for optimal dilution of sera and concentration of the engineered ACE2. DMS for infectivity and escape was performed in the setting of ∼3% infected cells among ∼15% spike-expressing cells.

**Fig. S6.**
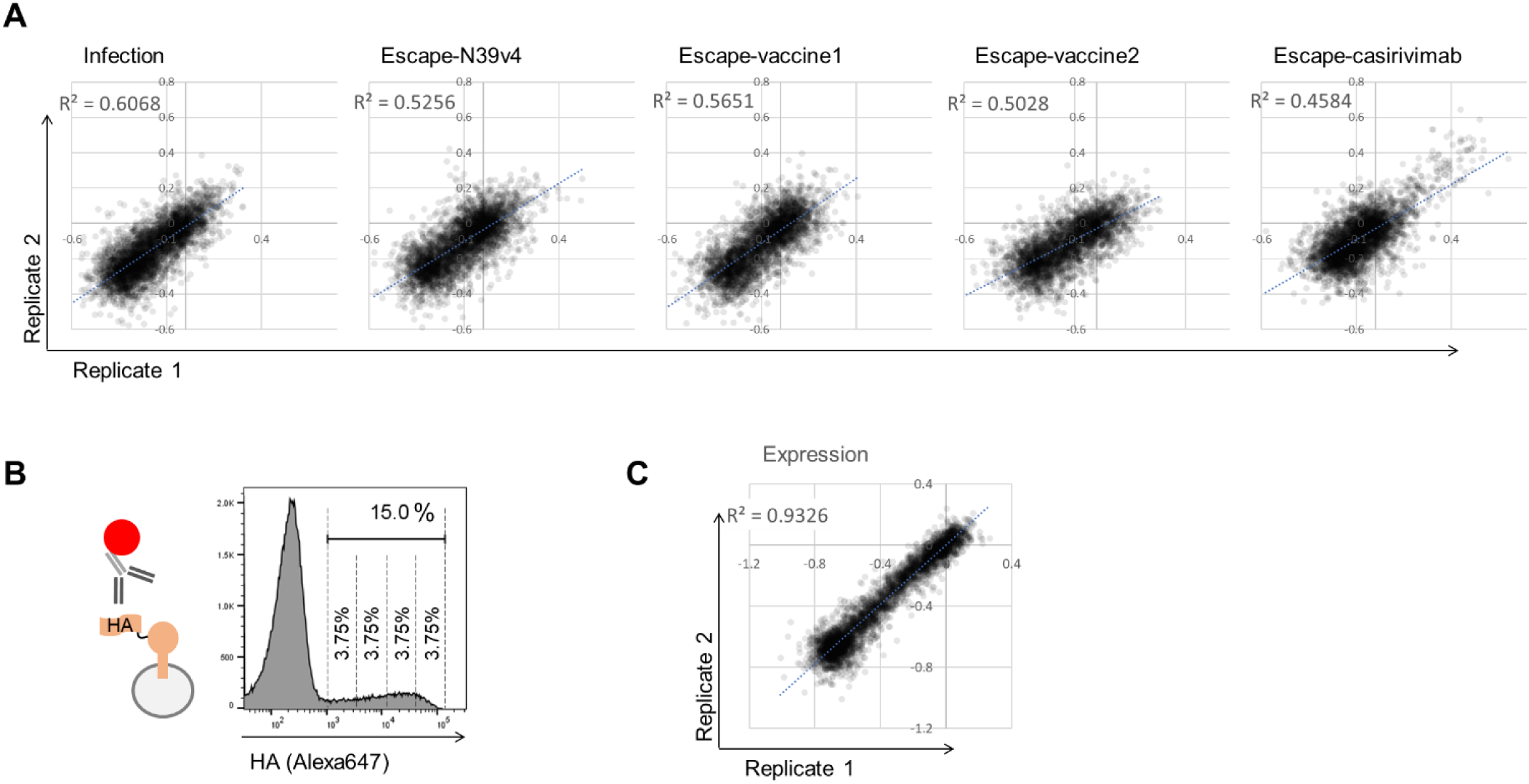
Reproducibility of each deep Mutational Scan for infectivity and escapes from neutralizing agents. (**A**) Correlation in mutation effects on the alteration of infectivity or escape from each neutralizing agent across replicates. (**B**) The gating of FACS to perform DMS for spike expression based on the staining of HA tag of spike library. Among ∼15% HA positive cells, the top 25% and bottom 25% of cells were sorted. (**C**) Correlation in mutation effects on the alteration of spike expression across replicates.

**Fig. S7.**
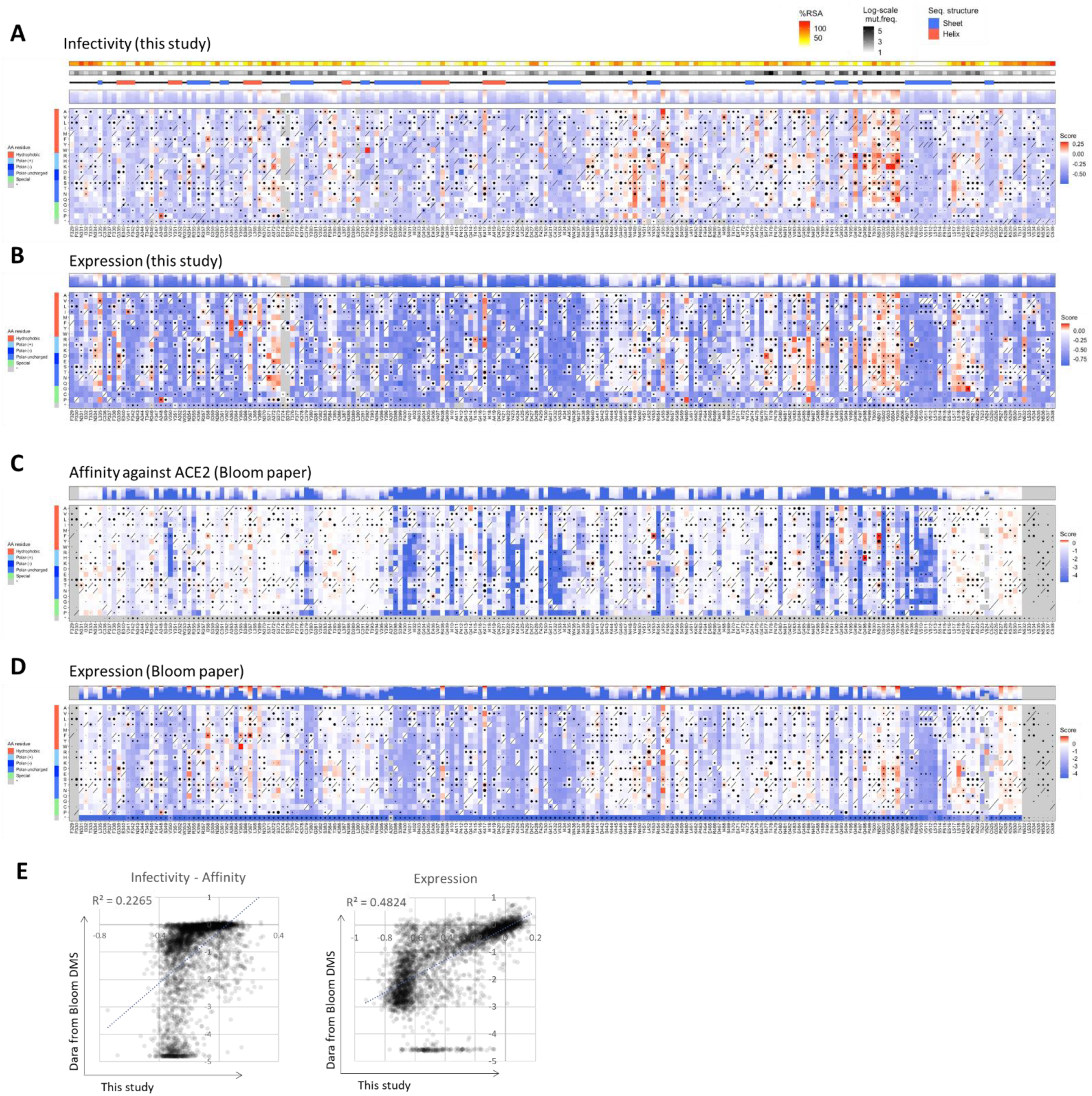
Sequence-to-Phenotype Maps of the whole RBD for infectivity and expression. (**A, B**) Heatmaps illuminating how all single mutations affects infectivity (A) and spike expression (B) in this study. (**C, D**) Heatmaps of DMS retrieved from Bloom paper (ref 18) for affinity against ACE2 (C) and RBD expression (D). (**E**) Scatter plot showing the correlation between this study and Bloom DMS. Left panel compares infectivity of this study and affinity of Bloom DMS. Right panel is expression level. Squares are colored by mutational effect according to scale bars on the right, with blue indicating deleterious mutations. Black dot size reflects the frequency in the virus genome sequence according to GISAID database as 17^th^ Dec 2021. %RSA is the relative solvent accessibility in the crystal structure of ACE2-Spike complex.

**Fig. S8.**
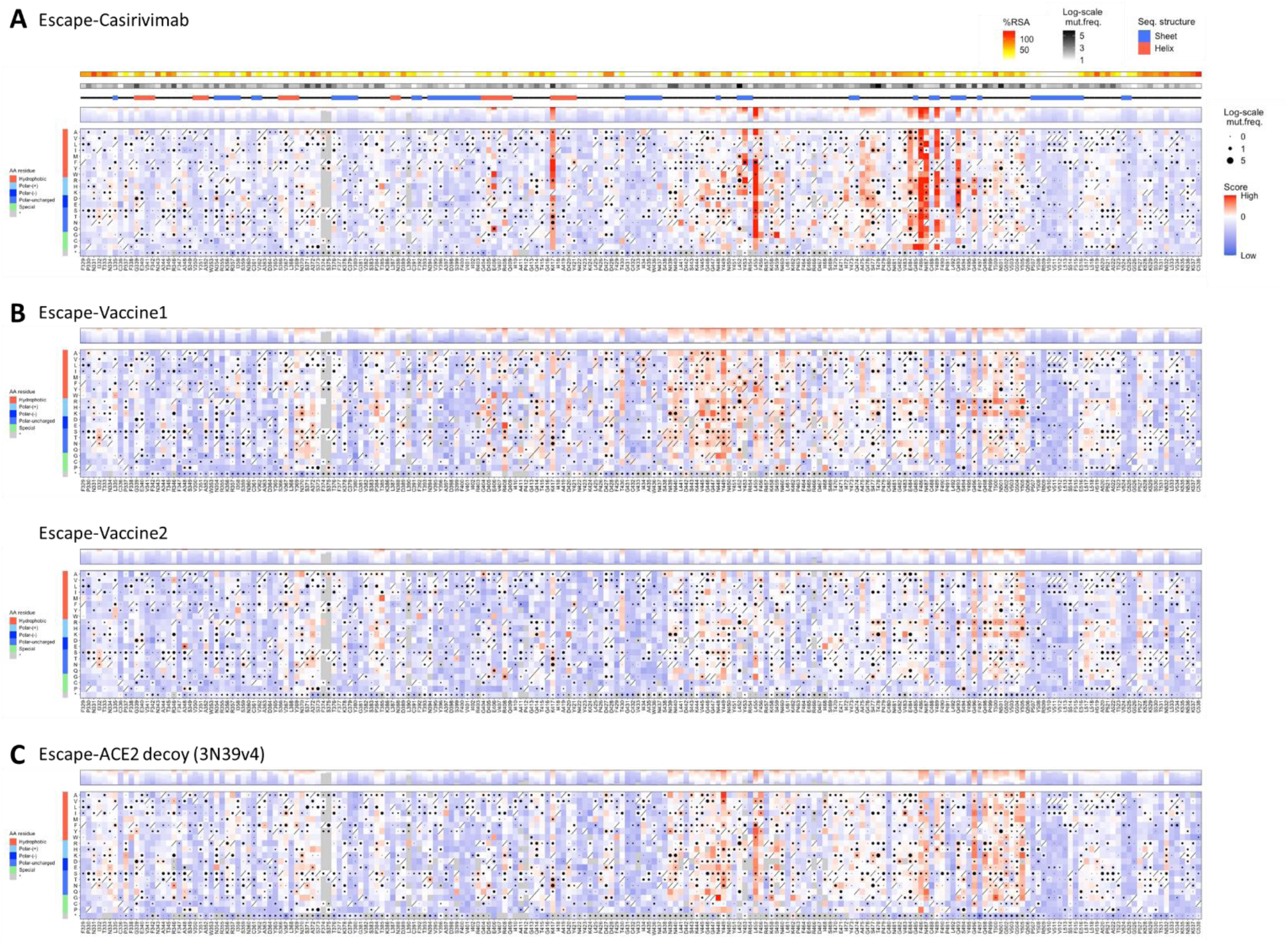
Sequence-to-Phenotype Maps of the whole RBD for escape. (**A**-**D**) Heatmaps illuminating how all single mutations affects infectivity (A), escape from casirivimab (B), vaccinated sera (C) and engineered ACE2 (D). Squares are colored by mutational effect according to scale bars on the right, with blue indicating deleterious mutations. Black dot size reflects the frequency in the virus genome sequence according to GISAID database as 17^th^ Dec 2021. %RSA is the relative solvent accessibility in the crystal structure of ACE2-Spike complex.

**Fig. S9.**
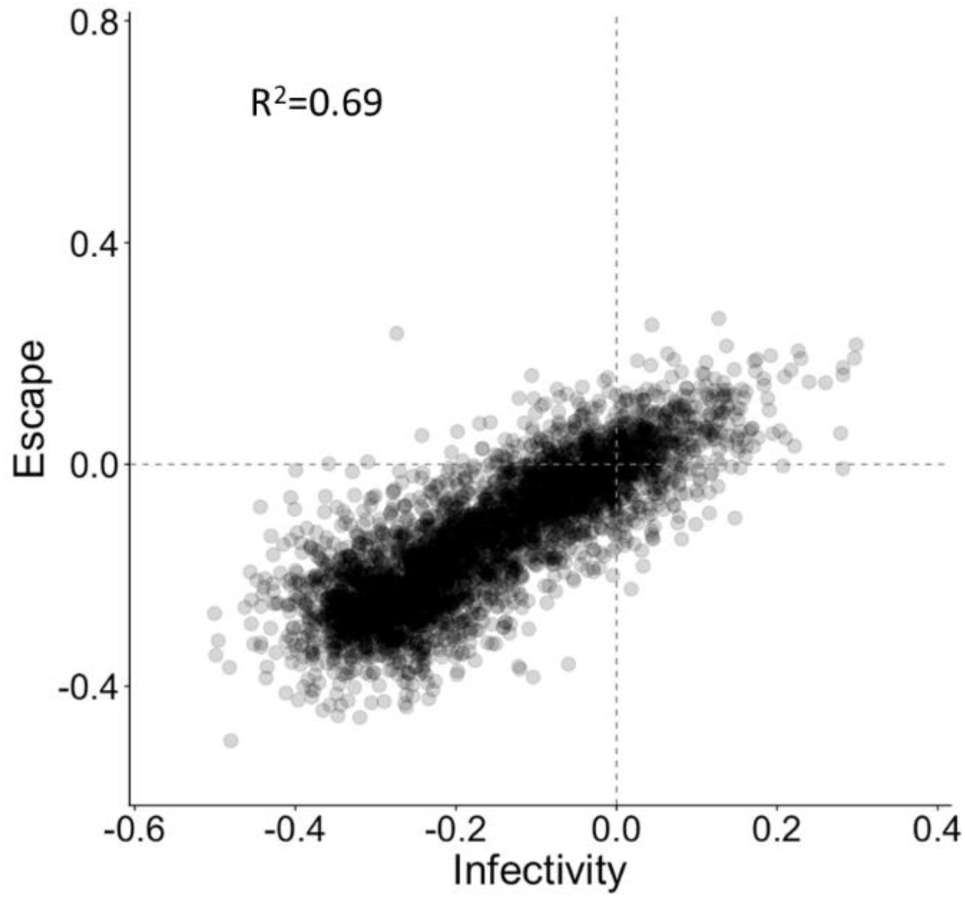
Reproducibility of correlation between infectivity and escape in the vaccinated sera. Correlation in mutation effects on the alteration of infectivity or escape from the neutralization by sera from different vaccinated individual.

